# Microbial interactions in the mosquito gut determine *Serratia* colonization and blood feeding propensity

**DOI:** 10.1101/2020.04.14.039701

**Authors:** Elena V. Kozlova, Shivanand Hegde, Christopher M. Roundy, George Golovko, Miguel A. Saldaña, Charles E. Hart, Enyia R Anderson, Emily A Hornett, Kamil Khanipov, Vsevolod L. Popov, Maria Pimenova, Yiyang Zhou, Yuriy Fovanov, Scott C. Weaver, Andrew L. Routh, Eva Heinz, Grant L. Hughes

**Affiliations:** Department of Pathology, University of Texas Medical Branch, Galveston, TX, USA; Departments of Vector Biology and Tropical Disease Biology, Centre for Neglected Tropical Disease, Liverpool School of Tropical Medicine, Liverpool, UK; World Reference Center for Emerging Viruses and Arboviruses, Institute for Human Infections and Immunity, and Department of Microbiology and Immunology, University of Texas Medical Branch, Galveston, TX, USA; Department of Pharmacology and Toxicology, Sealy Center for Structural Biology, University of Texas Medical Branch, Galveston, TX, United States; Department of Paediatrics and Tropical Medicine, Baylor College of Medicine, Houston, TX, United States; The Institute for Translational Science, University of Texas Medical Branch, Galveston, TX, United States. Institute for Global Health and Translational Science and SUNY Center for Environmental Health and Medicine, SUNY Upstate Medical University, Syracuse, NY, United States; Institute of Integrative Biology, University of Liverpool, Liverpool, UK; Department of Biochemistry and Molecular Biology, University of Texas Medical Branch, Galveston, TX, USA; Departments of Vector Biology and Clinical Sciences, Liverpool School of Tropical Medicine, Liverpool, UK

## Abstract

How microbe-microbe interactions dictate microbial complexity in the mosquito gut is unclear. Previously we found that *Serratia*, a gut symbiont that alters vector competence and is being considered for vector control, poorly colonized *Aedes aegypti* yet was abundant in *Culex quinquefasciatus* reared under identical conditions. To investigate the incompatibility between *Serratia* and *Ae. aegypti*, we characterized two distinct strains of *Serratia marcescens* from *Cx. quinquefasciatus* and examined their ability to infect *Ae. aegypti*. Both *Serratia* strains poorly infected *Ae. aegypti*, but when microbiome homeostasis was disrupted, the prevalence and titers of *Serratia* were similar to the infection in its native host. Examination of multiple genetically diverse *Ae. aegypti* lines found microbial interference to *S. marcescens* was commonplace, however one line of *Ae. aegypti* was susceptible to infection. Microbiome analysis of resistant and susceptible lines indicated an inverse correlation between *Enterobacteriaceae* bacteria and *Serratia*, and experimental co-infections in a gnotobiotic system recapitulated the interference phenotype. Furthermore, we observed an effect on host behaviour; *Serratia* exposure to *Ae. aegypti* disrupted their feeding behaviour, and this phenotype was also reliant on interactions with their native microbiota. Our work highlights the complexity of host-microbe interactions and provides evidence that microbial interactions influence mosquito behaviour.

## Introduction

Mosquitoes harbour a variety of diverse microbes that profoundly alter host phenotypes^1-3^. In general, the bacterial microbiome can vary considerably between mosquito species and individuals, but within an individual, it is comprised of relatively few bacterial taxa^4,5^. It is becoming more apparent that a variety of factors contribute to this variation, but we have a lack of understanding regarding why some taxa are present in a host, yet others are absent. In mosquitoes and other insects, much effort has been undertaken to characterize the infection status of species and populations for specific endosymbiotic bacteria such as *Wolbachia*^6-9^, yet few studies have examined the infection prevalence of specific gut-associated bacteria in mosquito vectors. It is evident that several gut-associated bacterial taxa are common between phylogenetically diverse mosquito species^4,5^, but less attention has been paid to identifying incompatible host-microbe associations and the mechanism(s) behind this incompatibility.

Microbiome assembly in mosquitoes is influenced by the environment, host and bacterial genetics, and stochastic processes. While the host is instrumental in maintaining microbiome homeostasis^10-14^, evidence is emerging that bacterial genetics and microbe-microbe interactions also dictate the prevalence and abundance of microbiota^15-18^. It is clear that the microbiome can influence several important phenotypes in mosquito vectors^3,19,20^, including the ability to transmit pathogens. Therefore, a greater appreciation of factors that influence colonization of the mosquito gut could assist our understanding of mosquito phenotypes important for vectorial capacity. This will be critical for deploying microbial-based approaches to control mosquito-borne disease^21,22^.

*Serratia* is a ubiquitous genus of gut symbionts that is known to infect a diverse array of insects, including taxa within the Homopteran, Hymenopteran, Dipteran, and Lepidopteran orders^23-28^. Several medically relevant vectors also harbour this bacterium^29-33^. In mosquitoes, *Serratia* appears to broadly infect Culicine and *Anopheles* mosquitoes^34-37^, and these infections can have important phenotypic effects including altering the ability of these vectors to transmit pathogens^38-42^. Intriguingly, after sequencing the microbiome of *Culex quinquefasciatus, Ae. albopictus* and *Ae. aegypti* mosquitoes reared under identical conditions within the same insectary, we found a species-specific infection cline in *Serratia* levels^4^. *Serratia* was a dominant member of the microbiota within *Cx. quinquefasciatus*, infected *Ae. albopictus* at low levels, and poorly infected or was absent from *Ae. aegypti*^4^. We also found that field-collected *Ae. aegypti* from the Houston region (USA) lacked *Serratia*^4^.

Other studies have found variable results in respect to the prevalence of *Serratia* in the yellow fever mosquito. Using high throughput 16S rRNA amplicon sequencing, *Serratia* was found to be absent or at low levels in some *Ae. aegypti* field populations^5,38-41^, yet present in others^35,42^. Culture dependent approaches have also confirmed the presence of *Serratia* in this mosquito species^43-46^. The variable nature of infection in the field could be due to the presence or absence of this bacterium in the local aquatic environment; however, this does not explain the infection cline we observed in our insectary when rearing *Ae. aegypti* given that *Cx. quinquefasciatus* reared in the same insectary was heavily infected^4^. The lack of *Serratia* infection in these lab-reared *Ae. aegypti* mosquitoes suggests there is a maladaptation between this particular mosquito line and *Serratia* strains.

To investigate the incompatibility between *Serratia* and the yellow fever mosquito, we isolated and characterized two distinct strains of *S. marcescens* present within the *Cx. quinquefasciatus* microbiome, and examined their ability to infect *Ae. aegypti*. We found that both *S. marcescens* strains poorly infected several *Ae. aegypti* strains. However, inducing dysbiosis in the native microbiota with antibiotics facilitated infections, suggesting the incompatibility was related to microbe-microbe interactions. In addition to microbial antagonism, we found that infection with these *S. marcescens* strains disrupted the feeding behaviour of mosquitoes. We further show the phenotypes induced by *S. marcescens* are driven by interactions with *Enterobacteriaceae* bacteria. Our work highlights the complexity of host-microbe interactions and provides further evidence that microbial exclusion influences microbiome composition and abundance within mosquitoes. These results are also relevant in the context of the holobiont, whereby both the host and the associated microbiota dictate organism phenotypes.

## Methods

### Mosquito rearing

Colony mosquitoes were reared at 27°C with 80% humidity in the UTMB insectary. Mosquitoes were fed 10% sucrose ad libitum and maintained at a 12:12 light:dark cycle. Mosquitoes were fed with defibrinated sheep blood (Colorado Serum Company) using a hemotek membrane feeder. **Table S1** lists the colony mosquitoes used in experiments.

### Isolation and characterisation of *S. marcescens* from *Culex quinquefasciatus*

Homogenates of *Cx. quinquefasciatus* were stored in PBS at *-*80°C as a glycerol stock. *S. marcescens* was isolated using conventional microbiological culturing. Briefly, LB plates were inoculated and incubated at 30°C. Individual bacterial colonies were selected and purified from two different *Culex* mosquitoes. Two *S. marcescens* strains, named CxSm1 and CxSm2, were selected. Both strains had a red pigmentation, although intensity of the colour varied between strains. Additinoally, there were differences in swimming motility and oxidase activity. These strains were sub-cultured for species identification by PCR amplifying the variable region of the 16S ribosomal RNA gene using universal bacterial primers. Primer sequences are listed in **Table S2**. Swimming motility was determined by inoculating LB medium (0.35% agar), incubating at 30°C overnight, and then quantifying motility toward the periphery of the plate^47^. DB BBL™ oxidase reagent droppers (BD & Comp., Sparks, MD) were used to detect cytochrome oxidase activity in bacteria following the manufacturer’s instructions. Scanning electron microscopy (SEM) was conducted as previously described^48,49^.

### Selection of *S. marcescens* antibiotic resistant mutants

*S. marcescens* antibiotic resistant mutants were created as described^50^ with some modification. Briefly, tubes containing 5 ml of LB broth with different concentrations of streptomycin (Sm) (Sigma) and rifampicin (Rif) (Sigma) (range: 0 [control] and 5 μg/ml, 10 μg/ml, 25 μg/ml, 50 μg/ml) were inoculated with 0.1 ml of a dilution of the bacterial cultures to obtain an inoculum of approximately 10^6^ CFU/ml. After overnight incubation at 30°C, bacterial aliquots from the tubes with the highest concentration of appropriated antibiotic were inoculated in LB broth [Sm supplemented (range: 0 [control], 25 μg/ml, 50 μg/ml, 100 μg/ml) and Rif supplemented (range: 0 [control], 50 μg/ml, 100 μg/ml, 200 μg/ml, 250 μg/ml, 500 μg/ml)] and incubated overnight at 30°C. Finally, after several passages in the presence of corresponding antibiotics, the CxSm1^RifR^ (MIC 400 μg/ml) and CxSm2^SmR^ (MIC 150 μg/ml) mutants were selected. The same approach was used to create a *Cedecea*^RifR^ mutant.

### Oral infection of mosquitoes with *S. marcescens*

The *S. marcescens* CxSm1^RifR^ and CxSm2^SmR^ strains were used for mosquito oral infection. Bacteria were grown in a 25 ml LB medium overnight culture at 30°C containing either Rif (200 μg/ml) or Sm (100 μg/ml). Bacteria were pelleted by centrifugation at 5000 rpm for 20 minutes and then washed twice with sterile PBS and suspended in 2.5 ml PBS. Bacterial PBS stock was titrated by serial dilutions and quantified by plating on LB agar and measuring colony forming units (CFUs). The bacterial PBS stock dilutions were resuspended in 10 % sterile sucrose to a final concentration of 1×10^7^ cells/ ml. When supplementing antibiotics in the sugar meal, Rif (200 μg/ml) or Sm (100 μg/ml) were added to the sucrose solution. Mosquitoes were fed with a bacterial infected solution for three days. Then, mosquitoes were fed with 10 % sterile sucrose or 10 % sterile sucrose plus corresponded antibiotic, as required. At each time point, ten mosquitoes from each group were aspirated, surface sterilized, and homogenized in 250 μl PBS separately. Serial dilutions of mosquito homogenate were plated on LB agar and LB agar with the appropriate antibiotic and CFUs quantified.

### Microbiome analysis of *Ae. aegypti* lines

The microbiomes of *Ae. aegypti* lines were analysed using barcoded high-throughput amplicon sequencing of the bacterial *16S rRNA* gene using a similar approach as previously described^4,51^. DNA was extracted (QIAamp DNA Mini kit) from individual whole surface sterilized mosquitoes five days post eclosion (N=15). To evaluative possible contamination, a spike in positive control^52^ was amplified under the same conditions as genomic DNA isolated from mosquitoes. The spike in control was synthesized as a gBlock (Intergrated DNA Technologies) and 100 pmole of template was used as template for PCRs. High-throughput sequencing of the bacterial 16S ribosomal RNA gene was performed using gDNA isolated from each sample. Sequencing libraries for each isolate were generated using universal 16S rRNA V3-V4 region primers in accordance with Illumina 16S rRNA metagenomic sequencing library protocols^53^. The samples were barcoded for multiplexing using Nextera XT Index Kit v2. Sequencing was performed on an Illumina MiSeq instrument using a MiSeq Reagent Kit v2 (500-cycles). To identify the presence of known bacteria, sequences were analyzed using the CLC Genomics Workbench 11.0.1 Microbial Genomics Module. Reads were trimmed of sequencing adaptors and barcodes, and any sequences containing nucleotides below the quality threshold of 0.05 (using the modified Richard Mott algorithm) and those with two or more unknown nucleotides or sequencing adapters were removed. Reference based OTU picking was performed using the SILVA SSU v128 97% database^54^. Sequences present in more than one copy but not clustered to the database were placed into *de novo* OTUs (99% similarity) and aligned against the reference database with 80% similarity threshold to assign the “closest” taxonomical name where possible. Chimeras were removed from the dataset if the absolute crossover cost was three using a k-mer size of six. Alpha diversity was measured using Shannon entropy (OTU level), rarefaction sampling without replacement, and with 100,000 replicates at each point. Beta diversity was calculated and nMDS plots were created using Bray-Curtis dissimilarity. Differentially abundant bacteria (family level) were identified using analysis of composition of microbiomes (ANCOM) with a significance level of *p* < 0.05^55^.

The total *Serratia* load within each mosquito line was assessed by qPCR. The S-adenosylhomocysteine nucleosidase (PFS) gene of *Serratia* was amplified with the primers psf1-F and psf-R^56^. The *Ae. aegypti* or *Cx. quinquesfactius S7* gene was amplified with aeg-S7-F and aeg-S7-R or Cq-S7-F and Cq-S7-R primers respectively^4^. The PCR was done in a 10 μl reaction containing 1 μM of each primer, 1× SYBR Green (Applied Biosystems) and 2 μl of genomic DNA template. Cycling conditions involved an initial denaturation at 95°C for 10 min, 40 cycles of 15 s at 95°C, 1 min at 60°C. Fluorescence readings were taken at 60°C after each cycle before deriving a melting curve (60–95°C) to confirm the identity of the PCR product. The PCR was carried out on the ABI StepOnePlus Real-Time PCR System. Relative abundance was calculated by comparing the *Serratia* load to the single copy mosquito gene.

### Life history assays

To determine blood feeding success, mosquitoes were offered a sheep blood meal using a hemotek feeding system. Cups of 25 female mosquitoes were starved for 24 hours prior to blood feeding. Mosquitoes were given the opportunity to feed, and then the number of blood fed mosquitoes were counted. For a subset of mosquitoes, the prevalence of *S. marcescens* in blood-fed and non-blood fed mosquitoes was determined by plating on selective media. To examine the reproductive output, we measured the number of eggs produced by a blood feed female. Individual blood fed females were placed into a vial with an oviposition site. After 4 days, the number of eggs were counted. Females that did not lay were excluded from the analysis. For most assays, the mortality of mosquitoes was quantified daily by counting and removing dead mosquitoes in cups.

### Genome sequencing

DNA isolation from bacteria was done using the PureLink™ Genomic DNA Mini Kit (Thermo Scientific). The Oxford Nanopore Technologies’s (ONT) MinION libraries were created with 1D Native barcoding genomic DNA kit (with EXP-NBD103 and SQK-LSK108), following standard protocol (ver. NBE_9006_v103_revO_21Dec2016). In brief, 1.5 µg of each genomic DNA was fragmented (Covaris g-TUBE), end-repaired (NEBNext® Ultra™ II End Repair/dA-Tailing Module), barcodes are ligated, pooled in equal-molar amounts and finally adapter ligated. The pooled library was loaded to a FLO-MIN106 flow cell and sequenced using the default settings of the MinKNOW for at least 24 hours. Base-calling was conducted with *Albaco*re (release 2.3.3, https://nanoporetech.com/) with the following parameters: -k SQK-LSK108 -f FLO-MIN106 --barcoding. Data trimming and quality filtering was conducted with *Porechop* (https://github.com/rrwick/Porechop) with the following parameter: -- discard_unassigned.

In addition, bacterial strains were submitted for short-read Illumina sequencing to 30X coverage using the Standard Whole Genome Service from the MicrobesNG service (www.microbesng.com, Birmingham, UK). Assemblies were performed using unicycler^57^, generating a hybrid assembly using both long- and short-read sequences as input for each strain, respectively, for the assembly process. FastANI (average nucleotide identity) was used on a set of *Serratia* reference genomes retrieved from NCBI (**Table S3**) to confirm the species allocation. ANI analysis shows that CxSm1 and CxSm2 are highly similar to *Serratia* sp. Y25, which likely forms a subspecies of *S. marcescens* with an average ANI distance of 0.054 (**Table S4, Fig S1**); there was no difference in ANI level between CxSm1 and CxSm2. Mapping against *S. marcescens* reference strains thus resulted in high numbers of SNPs (252113 and 253191 for CxSm1 and CxSm2, respectively, against NZ_HG326223 DB11); whereas 44435 and 44913 SNPs were detected when mapping against the *Serratia* sp. YD25 genome (CP016948.1). 44169 of these were ACGT-only sites where at least one of the sequences differs from the reference; 29 of these core genome SNPs differ between CxSm1 and CxSm2. Mapping was performed using SMALT v0.7.4 (ref: SMALT: A mapper for DNA sequencing reads. Available from: https://sourceforge.net/projects/smalt/) to produce a BAM file. Variation detection was performed using SAMtools mpileup v0.1.19 (PMID:19505943) with parameters “-d 1000 -DSugBf” and bcftools v0.1.19 (ref: bcftools: Utilities for variant calling and manipulating VCFs and BCFs. Available from: http://samtools.github.io/bcftools/) to produce a BCF file of all variant sites. The option to call genotypes at variant sites was passed to the bcftools call. All bases were filtered to remove those with uncertainty in the base call. The bcftools variant quality score was required to be greater than 50 (quality <□50) and mapping quality greater than 30 (map_quality <□30). If the same base call was not produced by all reads, the allele frequency, as calculated by bcftools, was required to be either 0 for bases called the same as the reference, or 1 for bases called as a SNP (af1□<□0.95). The majority base call was required to be present in at least 75% of reads mapping at the base (ratio□<□0.75), and the minimum mapping depth required was 4 reads, at least two of which had to map to each strand (depth <□4, depth_strand <□2). Finally, strand_bias was required to be less than 0.001, map_bias less than 0.001, and tail_bias less than 0.001. If any of these filters were not met, the base was called as uncertain. The raw sequence data have been deposited to SRA/ENA and are available under accessions [awaiting accession numbers], the assemblies are available from GenBank under accessions [awaiting accession numbers].

## Results

### *Serratia* strain characterization

Two strains of *Serratia* were isolated from *Cx. quinquefasciatus* by conventional microbiology procedures. 16S rRNA sequencing indicated these strains were *S. marcescens*, and each produced a red pigmentation when grown in a culture which is indicative of this species **(Fig S2A)**. Although the 16S rRNA sequence was identical between strains, we saw phenotypic differences in their swimming motility, oxidase activity, and capacity to form biofilms, suggesting they were phenotypically divergent (**Fig S2A-B**). Swimming motility has been implicated in host gut colonization of several hosts^58,59^, and these traits can influence pathogen infection in mosquitoes^60^. To further characterize these strains (named CxSm1 and CxSm2), we sequenced their genomes using nanopore and Illumina technologies. Comparative genome analysis indicated high similarity between the two strains, and both showed 94.7% average nucleotide identity (ANI) similarity to *S. marcescens* when comparing with a set of *Serratia* reference genomes, indicating that these might represent a subspecies of *S. marcescens* **(Fig S1, Table S3 and S4).** Recent work has indicated a population structure in *S. marcescens* with at least two different clades^61^, which might be an indication for several subspecies or indeed a species complex, as is for example seen for *Klebsiella pneumoniae* or *Enterobacter cloacae*^62,63^.To aid our recovery of each of these *S. marcescens* strains on media, we selected for rifampicin and streptomycin spontaneous antibiotic resistant isolates for CxSm1 and CxSm2 respectively (antibiotic resistant strains named CxSm1^RifR^ and CxSm2^SmR^).

### *Serratia* colonization of mosquitoes

We investigated the ability of *Serratia* to colonize the novel *Ae. aegypti* host by reinfecting bacteria into mosquitoes in a 10% sucrose meal and monitored infection dynamics in the mosquito over time. The *Serratia* infection was completely lost from *Ae. aegypti* by 12 dpi, whereas the bacterial prevalence in the native host, *Cx. quinquefasciatus*, remained constantly high with infection levels ranging from 100% infection for CxSm1^RifR^ to 80% infection for CxSm2^SmR^ at 12dpi (**Fig 1A**). Of the mosquitoes that were infected, both *S. marcescens* strains infected *Ae. aegypti* (Galveston) at significantly lower densities compared to their native host, *Cx. quinquefasciatus* (**Fig 1A**). For example, at 3 dpi, we saw approximately 1000 times less *Serratia* in *Ae. aegypti* compared to *Cx. quinquefasciatus* (**Fig 1A**). We also examined other culturable microbiota by plating mosquito homogenates on non-selective LB plates, and in general, we saw few changes in the number of CFUs between groups in either *Ae. aegypti* or *Cx. quinquefasciatus* (**Fig S3**), suggesting *Serratia* infection had minimal effect on the total bacterial load of culturable microbiota in mosquitoes. The inability of *Serratia* to persistently infect *Ae. aegypti*, which was not observed for other bacteria (**Fig S3**), suggests that barriers, either of bacterial or host origin, were promoting the maladaption between these *Serratia* strains and this line of *Ae. aegypti*.

**Figure 1.**
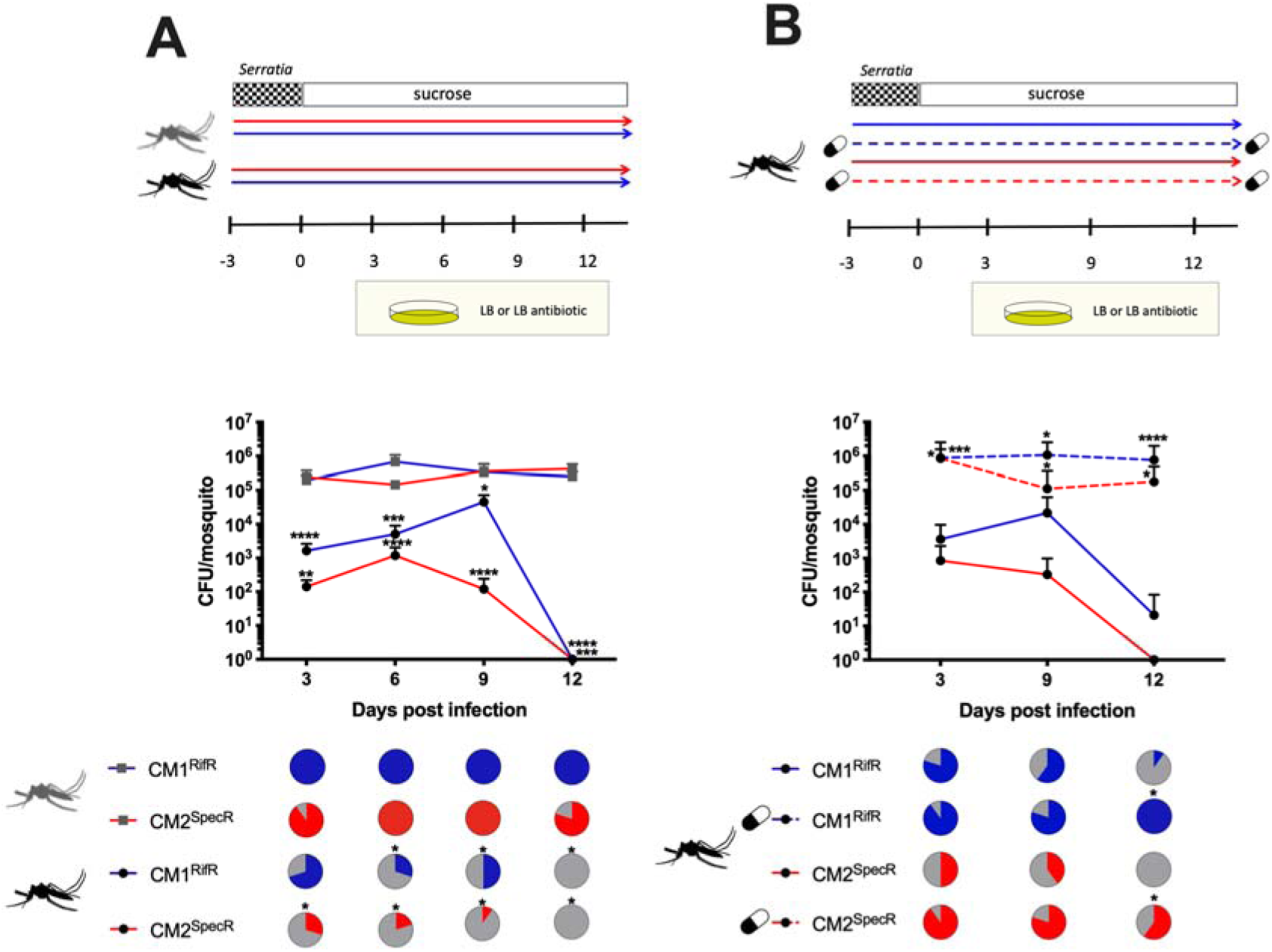
*Serratia* infections in native and non-native mosquito hosts. Infection of CxSm1^RifR^ and CxSm2^SmR^ into *Cx. quinquefasciatus* (grey) and *Ae. aegypti* (black) mosquitoes (**A**). Infection of CxSm1^RifR^ and CxSm2^SmR^ strains into antibiotic treated or untreated *Ae. aegypti* (**B**). Rifampamicin or spectinomycin was administered to mosquitoes in a sugar meal. For both A and B, the upper panel shows a schematic of experimental design. The line graph indicates the titer of *Serratia* in mosquitoes, and the pie graph indicates infection prevalence in mosquitoes. For each time point, ten mosquitoes were sampled. Letters indicate significance from Mann-Whitney test comparing density within a time point. Asterisks indicate a significant difference in *Serratia* in *Cx. quinquefasciatus* to *Ae. aegypti* (A) or antibiotic and non-antibiotic treated mosquitoes (B) using a Mann-Whitney test for densinty Fisher’s exact test for prevalence. * p <0.05, ** p < 0.01, *** p < 0.001, **** p < 0.0001.

### Microbial intereaction in the mosquito gut

To gain insights into the mechanism promoting the incompatibility between *Serratia* and *Ae. aegypti*, we repeated infections in antibiotic treated mosquitoes as we speculated that the native microbiota of mosquitoes might interfere with the colonization of the host (**Fig 1B**). We formulated this hypothesis as we have previously seen evidence of bacterial exclusion of symbiotic microbes in mosquitoes^4,18^. Strikingly, both CxSm1^RifR^ and CxSm2^SmR^ colonized mosquitoes at significantly higher titers when mosquitoes were treated with antibiotics compared to mosquitoes reared conventionally without antibiotics (Mann Whitney test; CxSm1^RifR^; day 3 p < 0.002, day 9 p < 0.01; day 12 p < 0.0001, CxSm2^SmR^; day 3 p < 0.03, day 9 p < 0.01; day 12 p < 0.01) (**Fig 1B**). Furthermore, for both *Serratia* strains, significantly more mosquitoes were infected at day 12 in antibiotic treated mosquitoes compared to untreated (Fisher’s exact test; CxSm1^RifR^ p = 0.01, CxSm2^SmR^ p = 0.0007). The levels of *Serratia* in *Ae. aegypti* after microbiome homeostasis was disrupted by antibiotics were comparable to infections in the native host *Cx. quinquesfasciatus* (**Fig 1A**). These data indicated that the *Ae. aegypti* (Galveston) line had the capacity to harbor *Serratia*, and that the incompatibility in mosquitoes with an intact microbiome (**Fig 1A**,^4^) was due to members of the native microbiota inhibiting *Serratia*, as opposed to intrinsic host factors or genetic factors of the *S. marcescens* strains.

To determine how widespread these microbial interactions were in *Ae. aegypti* mosquitoes, we investigated eight diverse lines for native *Serratia* infections and their capacity to be infected with CxSm1^RifR^. When examining the native *Serratia* load by qPCR, seven of the eight *Ae. aegypti* lines had significantly lower titers compared to *C. quinquefasciatus* (**Fig. 2A**). Intriguingly, an *Ae. aegypti* line from Thailand had a high *Serratia* load that was comparable to the infection in the native *Culex* host. We also quantified *Serratia* levels in two other *Cx. quinquefasciatus* lines and found similar or higher loads of *Serratia* in these other lines (**Fig S4**), indicating the robust infection of *Serratia* in *Cx. quinquefasciatus* was commonplace. We then infected the CxSm1^RifR^ *Serratia* strain into these eight diverse *Ae. aegypti* lines. For these infections we focused our attention on CxSm1^RifR^, as overall, it appears this strain had a greater capacity to infect *Ae. aegypti* compared to CxSm2^SmR^. We therefore posited that this strain would be more likely to infect non-native hosts. Similar to our previous experiments, *Serratia* poorly infected the Galveston line and was eliminated by 12 dpi. In the other lines, we saw some variation in the time it took for *Serratia* to be eliminated, with clearance occurring rapidly in the Juchitan and Iquitos lines. Whilst the process took longer in others (Dakar, Salvador and Dominican Republic), infection was ultimately cleared from all lines. In stark contrast to these seven lines, the Thailand line harboured the *Serratia* infection at similar levels compared to the native *Culex* host. Combined, the qPCR and re-infection experiments indicated the majority of *Ae. aegypti* lines were not permissive to *Serratia* infection, but infection dynamics in the Thailand line were similar to the native *Culex* host.

**Figure 2.**
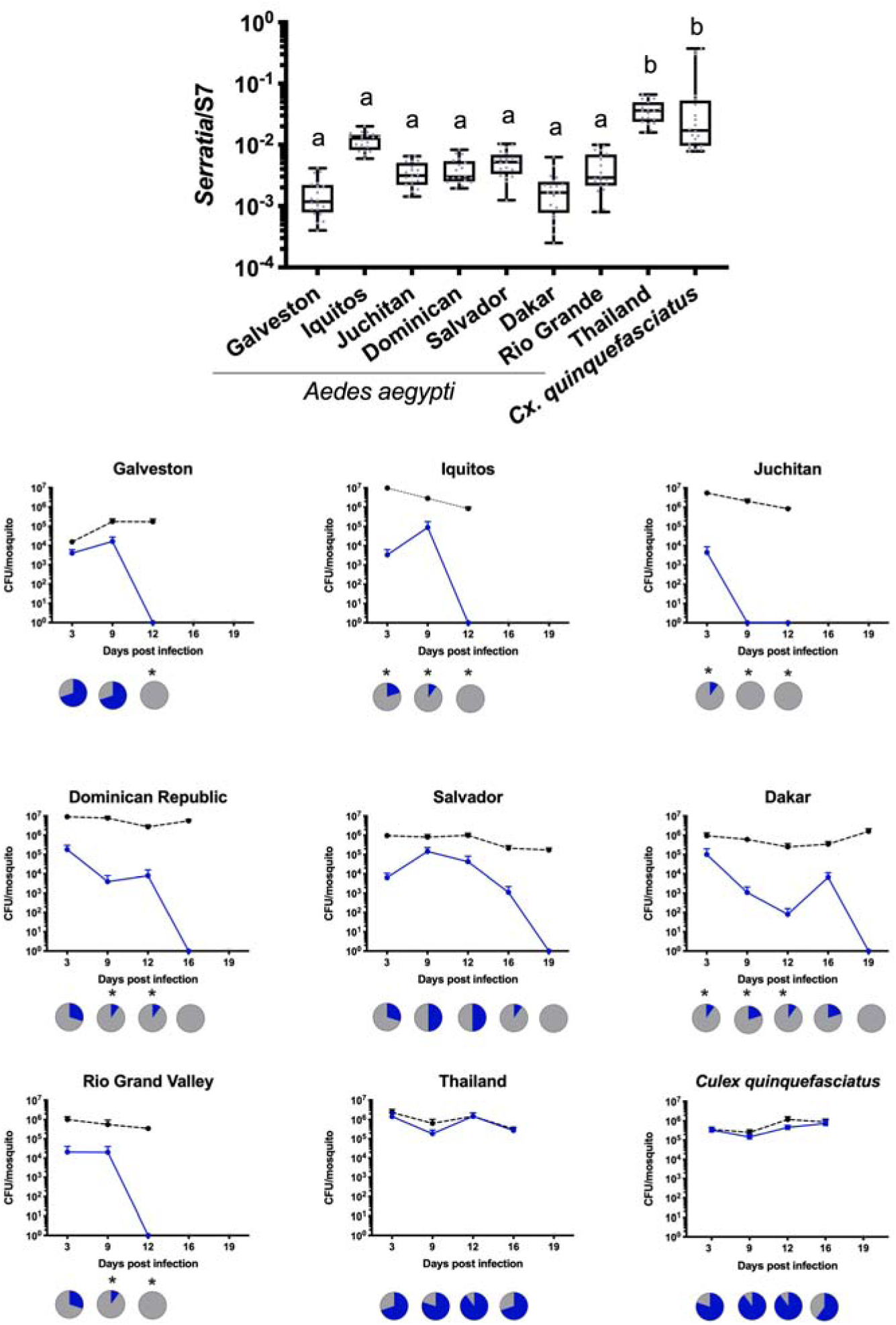
*Serratia* infection in diverse *Ae. aegypti* lines. The density of *Serratia* was determined by qPCR in eight *Ae. aegypti* lines (**A**). CxSm1^RifR^ was infected into Ae. aegypti lines and density (blue line) monitored over time (**B**). Total cultural microbiota (dotted line) was also quantified by culturing bacteria from homogenized mosquitoes on non-selective LB plates. Line graphs indicate titer of *Serratia* in mosquitoes, and pie graphs indicate infection prevalence in mosquitoes. Asterisks indicate a significant difference in *Serratia* prevalence in *Ae. aegypti* lines compared to *Cx. quinquefasciatus* using a Fisher’s exact test compared to the *Cx. quinquefasciatus* control line.

### Microbiome analysis of resistant and susceptible mosquitoes

To determine which specific microbiota of *Ae. aegypti* altered *Serratia* infections, we sequenced the microbiome of four select lines that varied in the capacity to harbor the bacterium. The V3-V4 variable region of the 16S rRNA gene was sequenced from the Juchitan, Galveston, and Iquitos line, which were recalcitrant to *Serratia*, and the Thailand line which was able to sustain the infection similar to the native host. For each line, we sequenced 15 individuals and, on average, obtained 32,000 reads per mosquito. Rarefaction curves indicated that sufficient depth was obtained in the sequencing to adequately characterize the microbiome, while our spike in controls constituted 99.5% of the relative abundance indicating that there was negligible contamination in our sequencing (**Fig S5**). Across all mosquito lines, we identify a total of 1,163 bacterial OTUs, but only 55 were present in mosquitoes at an infection frequency above 1% (**Table S5)**.

When examining taxa within the microbiome, the majority of sequences were from the Proteobacteria, while others were associated with Verrucomicrobia and Bacteroidetes. Within the Proteobacteria, the most abundant OTUs were in with the families *Enterobacteriaceae, Acetobacteriaceae, and Pseudomonasaceae*, while the Thailand line harboured a considerable amount of *Verrucomicrobiaceae* compared to the other three lines **(Fig 3A)**. Confirming our qPCR data, we saw minimal or no *Serratia* infection in the Galveston, Iquitos, or Juchitan lines, but this bacterium comprised approximately 4% of the relative abundance of the Thailand line (**Fig 3B**). It was also noticeable that the Thailand line possessed many more OTUs compared to the other lines (**Fig S6; Table S5**). This was corroborated by alpha diversity measures, which indicated the Thailand line had a significantly elevated Shannon’s diversity index compared to the other three lines (**Fig 3C**). To examine the community structure of the microbiome in each line, we undertook non-metric multidimensional scaling (NMDS) analysis based on Bray-Curtis dissimilarity. Strikingly, the microbiomes of each line were significantly different from one other (**Fig 3D**, p < 0.05), however it was evident from the pattern of clustering of the microbiota, that the Thailand line was considerably divergent compared to the other three lines.

**Figure 3.**
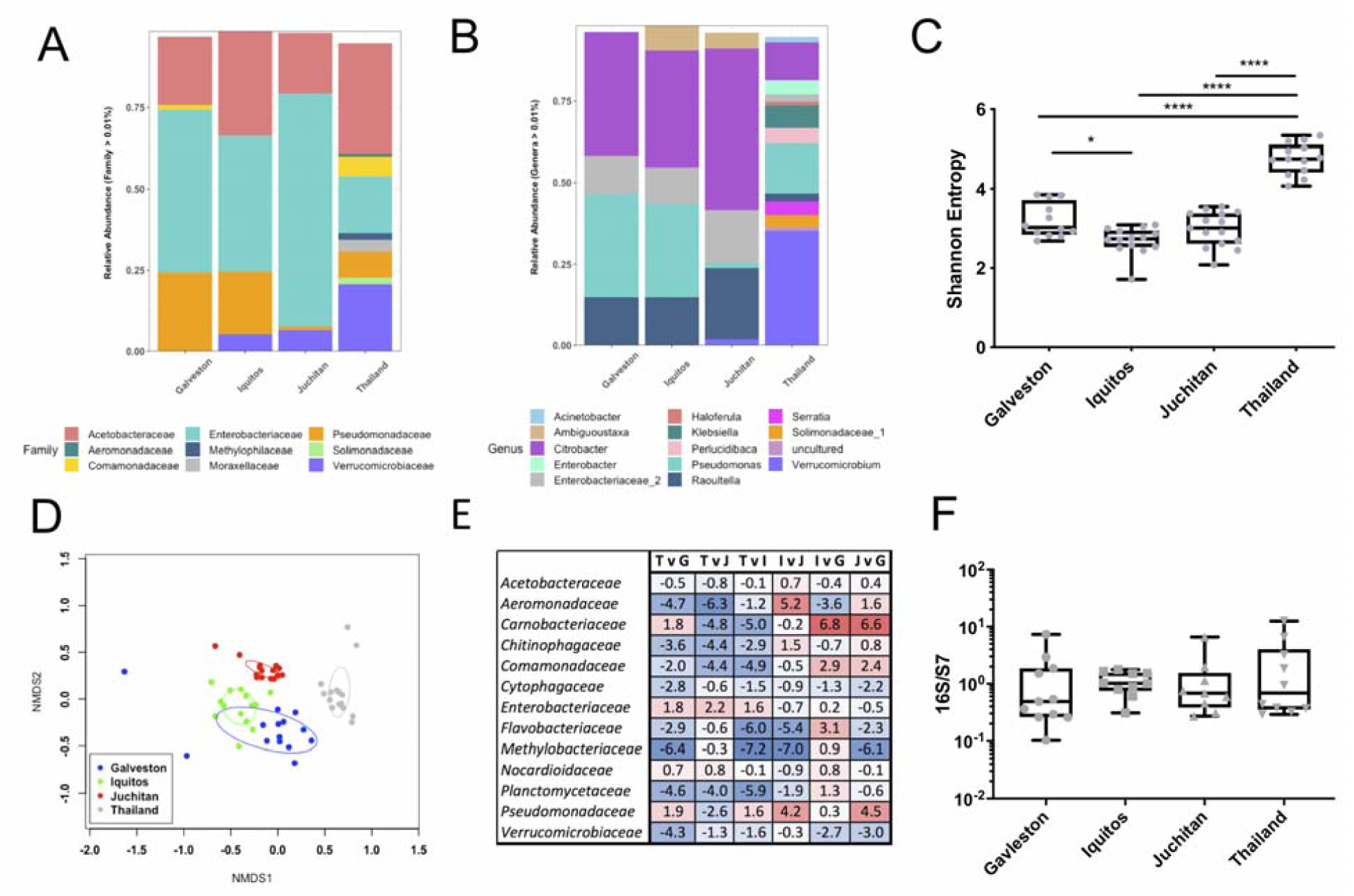
Microbiome analysis of the Galveston, Juchitan, Iquitos, and Thailand *Ae. aegypti* lines. 16S rRNA amplicon sequencing was done on female adult mosquitoes 5 days post eclosion. All mosquitoes were reared in the same laboratory environment under identical conditions. The relative abundance of bacterial communities at the family (**A**) and genus level **(B**). Alpha (Shannon’s entropy; * p <0.05, **** p < 0.0001) (**C**) and beta (NMDS) **D**) diversity metrics. Differential abundance analysis (ANCOM) of bacterial families in pairwise comparisons **(E)** between the four lines (T – Thailand, G – Galveson, J – Juchitan, I – Iquitos). A bolded value indicates a significant difference. Positive value indicates greater adundance of bacteria in the denominator, negative indicates greater number of bacteria in the numerator in the pairwise comparison. Total bacterial load in mosquito lines measured by qPCR **(F).**

To examine specific taxa that may be the cause of microbial incompatibility, we undertook pairwise comparisons to identify bacteria that were differentially abundant between lines. We examined differences at the family level using ANCOM, which is specifically designed to handle variable microbiome data^55^. While the abundance of several families was significantly different between lines, the *Enterobacteriaceae* were the only family that was consistently reduced in the Thailand line compared to the other three lines (**Fig 3E**). In addition to amplicon sequencing, we used qPCR to determine the total microbial load of mosquitoes and found each possessed a similar density of bacteria (**Fig 3F**), indicating the increase in taxa in the Thailand line were not simply due to possessing a greater number of bacteria. Taken together, these data indicated that the microbiome of the Thailand line was substantially different from the other lines and that members of the *Enterobacteriaceae* were candidate taxa that inhibited *Serratia* infection in mosquitoes.

### Co-infections in gnotobiotic infection model

To functionally demonstrate that *Enterobacteriaceae* interfered with *Serratia* colonization, we undertook a series of co-infection experiments in antibiotic treated mosquito lines. Prior to infection of the CxSm1^RifR^ *Serratia* strain, we infected mosquitoes with *Cedecea*^RifR^, an *Enterobacteriaceae* that commonly infects mosquitoes, or other *Acetobacteraceae* and *Pseuduomonadaceae* bacteria as controls (**Fig 4A**). We chose *Cedecea* as we have previously documented that this bacterium infects *Ae. aegypti* effectively^4^. The infection prevalence of *Serratia* in the co-infected *Ae. aegypti* Galveston line was significantly reduced in all time points (**Fig 4B**, p < 0.05, Fisher’s exact test). In the few mosquitoes that did harbour a *Serratia* infection, the density was significantly lower compared to the single infection (**Fig 4B**, t-test p < 0.05). These data indicated *Serratia* colonization was inhibited by the presence of *Cedecea*, and the phenotype we observed previously in conventionally reared mosquitoes could be recapitulated in a gnotobiotic setting. Similarly, we also found the prevalence of *Serratia* was reduced by co-infection in the *Ae. aegypti* Thailand line (p = 0.05, Fisher’s exact test), although this effect was more subtle, and no significant difference was observed at 12 dpi (**Fig 4C**). In contrast to co-infection with *Cedecea*, we found no effect in *Serratia* prevalence or titers when co-infected with *Asaia* or *Pseudomonas* (**Fig 4E and F**), which are members of the *Acetobacteraceae* and *Pseduomonadaceae* families, respectively. Interestingly, there was evidence that *Serratia* interferes with *Asaia* infections in *Ae. aegypti, as* there was an initial reduction in the prevalence of *Asaia* in the co-infected group compared to the single infection (**Fig 4F**, p = 0.05, Fisher’s exact test). Together, these co-infection studies demonstrate that inhibition of *Serratia* colonization in *Ae. aegypti* is bacteria specific, and that antagonism occurs between *Enterobacteriaceae* and *Serratia.*

**Figure 4.**
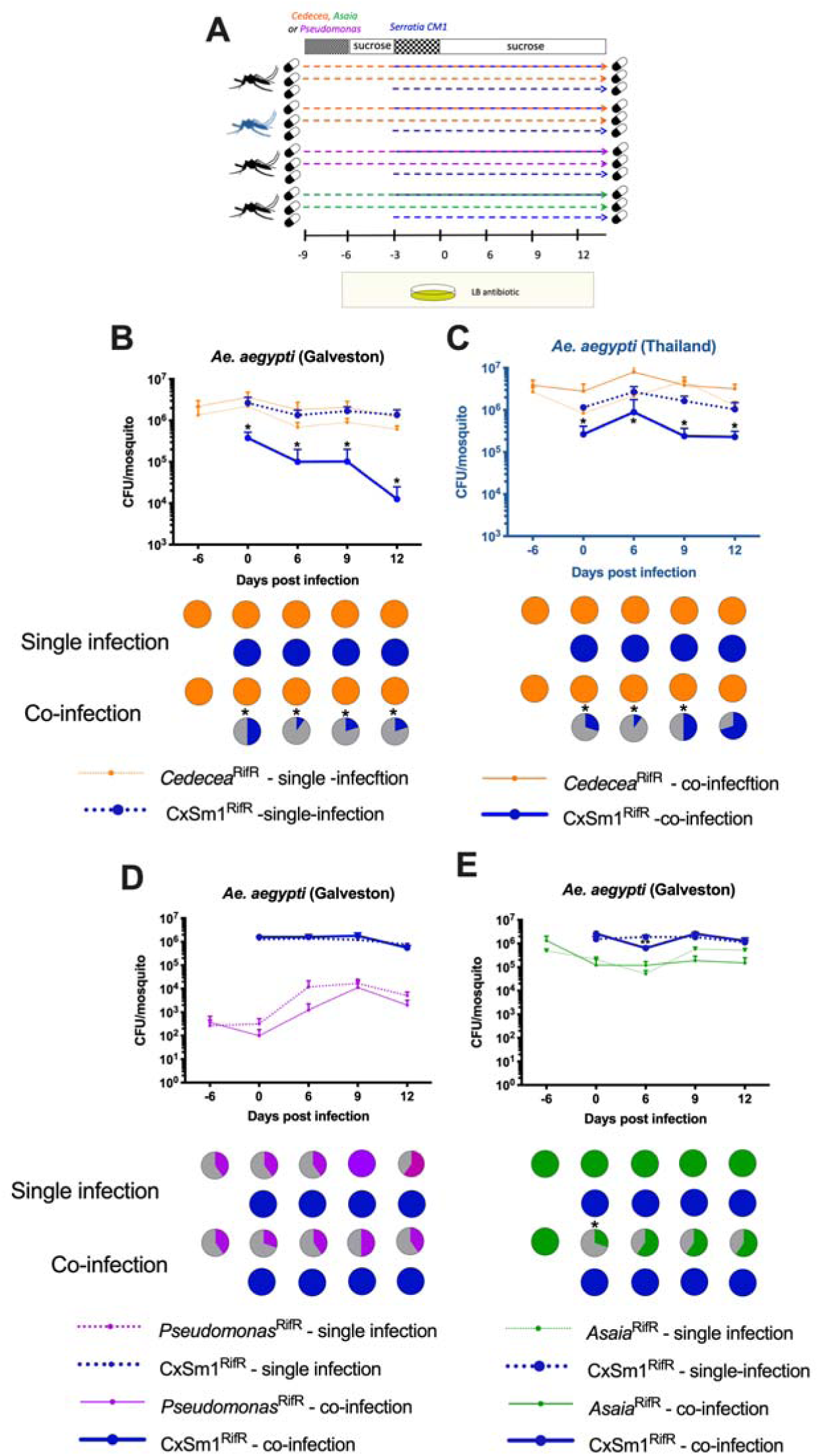
Co-infection of *Enterobacteriaceae* and *Serratia* in *Ae. aegpyti*. Schematic depicting the co-infection experimental design (**A**). Co-infection of *Cedecea*^RifR^ and CxSm1^RifR^ in *Ae. aegpyti* (Galveston) (**B**) and Thailand (**C**) lines. Control co-infections whereby *Pseudomonas* (**D**) or *Asaia* (**E**) were infected prior to CxSm1^RifR^. Line graphs show bacteria density (CFU/mosquito), and pie graphs show infection prevalence. For each time point, ten mosquitoes were sampled. Letters indicate significance from ANOVA comparing density within a time point. Asterisks indicate a significant difference between *Serratia* prevalence in single and co-infected mosquitoes using a Fisher’s exact test.

To determine how *Enterobacteriaceae* influenced *Serratia* in its native host, we repeated co-infection experiments in *Cx. quinquefasciatus* using the gnotobiotic infection model. *Cedecea* infected *Culex* mosquitos less effectively compared to *Aedes*, with infection densities around two logs lower and an infection prevalence that dropped to 50% over the course of the experiment (**Fig 5**). Despite a lower level of infection, *Cedecea* infection prior to *Serratia* reduced the infection of the latter. At 15 and 18dpi, the prevalence of *Serratia* in the co-infection was 50% compared to 100% in the single infection (**Fig 5A**, p = 0.03, Fisher’s exact test). We also examined the effect of *Cedecea* on an established *Serratia* infection by reversing the order each bacterium was administered to the mosquito. In this case, the prevalence of *Serratia* in the co-infection was significantly reduced only at the 18 dpi time point (**Fig 5B**, p = 0.03 Fishers exact test). Taken together, these data show that antagonism between *Serratia* and other *Enterobacteriaceae* also occurs in *Culex* mosquitoes.

**Figure 5.**
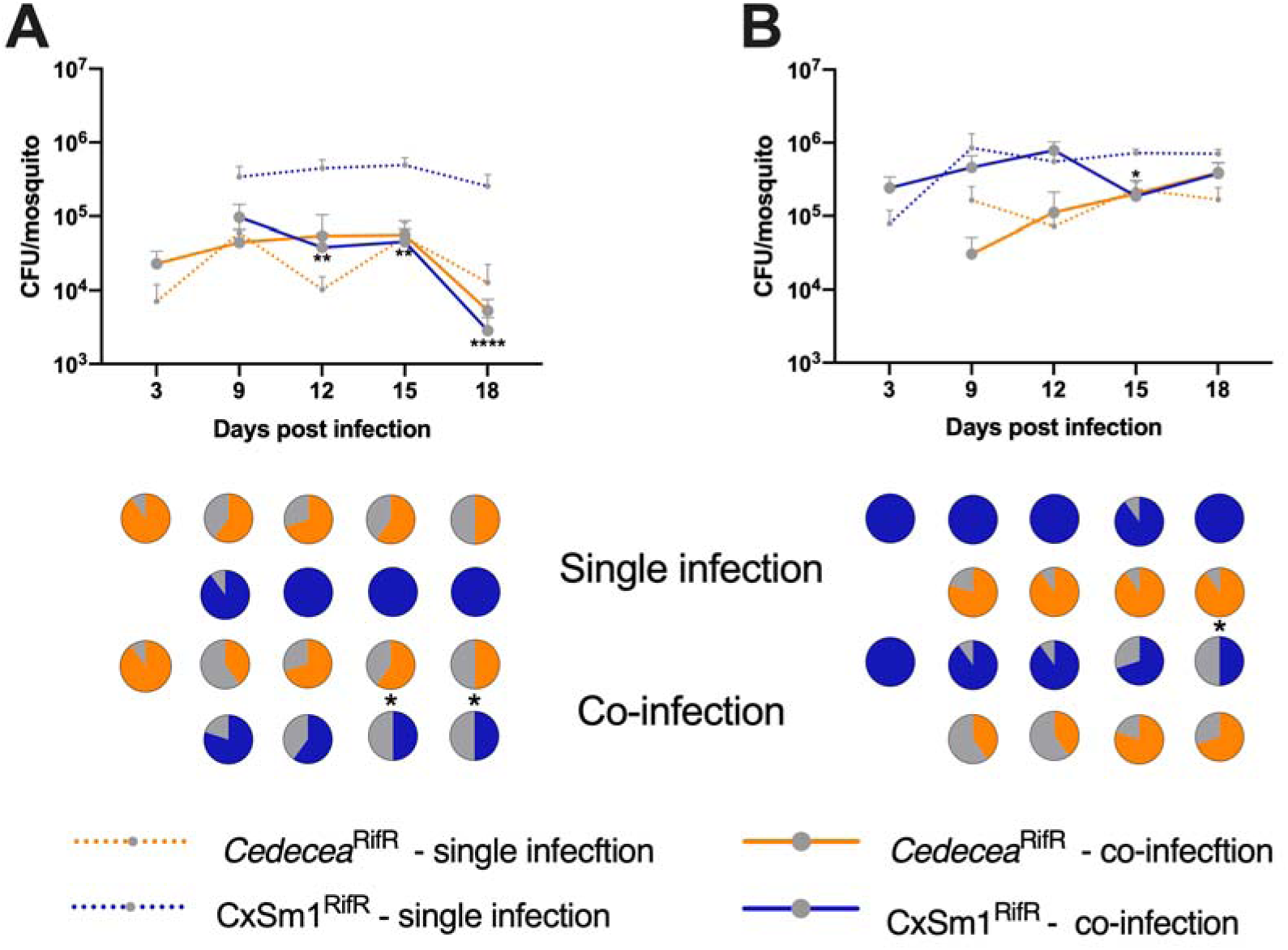
Co-infection of *Cedecea* and *Serratia* in *Cx. quinquefasciatus*. Infection of *Cedecea* followed by CxSm1^RifR^ (**A**) or CxSm1^RifR^ followed by *Cedecea* (**B**) in *Cx. quinquefasciatus.* Line graphs show bacteria density (CFU/mosquito), and pie graphs show infection prevalence. For each time point, ten mosquitoes were sampled. Letters indicate significance from Mann-Whitney test comparing density of CxSm1^RifR^ single and co-infectons within a time point. For prevalence data, asterisks indicate a significant difference between *Serratia* prevalence in single and co-infected mosquitoes using a Fisher’s exact test. * p <0.05, ** p < 0.01, **** p < 0.0001.

### Effect of *Serratia* exposure on blood feeding behaviour

Anautogenous mosquitoes require a blood meal to acquire nutrition for egg development. Ingested blood alters the gut microbiota composition and abundance, often increasing total bacterial load but decreasing species richness^64,65^. In other mosquito species, *Serratia* has been seen to rapidly increase in titer after a blood meal^66-68^ and, in some cases, can be lethal to the host^36^. As such, we investigated the influence of blood feeding on *Serratia* infected *Ae. aegypti* (**Fig 6A**). We measured bacterial load in the mosquito (**Fig 6B**) as well as a range of life history traits. For these experiments, we focused our attention on CxSm1^RifR^. In contrast to a previous study^36^, we observed no fitness costs to infection in terms of mosquito survival pre or post-blood meal (**Fig S7**). After a blood meal, *Serratia* density precipitously increased around 100-fold. The increase in the antibiotic treated mosquitoes was more subtle, likely because the bacterial load was initially greater, suggesting there is an upper limit to infections. After blood feeding, *Serratia* infections were comparable to densities and infection frequencies seen in sugar fed mosquitoes (**Fig 1 & 4**), with levels in antibiotic treated mosquitoes being maintained at around 1×10^6^ bacteria/mosquito. In conventionally reared mosquitoes, *Serratia* was eliminated, albeit over a longer time period, likely due to the increased density of the bacterium after stimulation from the blood meal. Post blood feeding, *Serratia* densities equilibrated to levels around 10^6^, which were comparable to infection densities seen in non-blood fed mosquitoes (**Fig 6B**). While we saw no differences in egg number (**Fig S8)**, in the process of conducting these experiments, we observed that CxSm1^RifR^ infected mosquitoes were less inclined to take a blood meal when reared on a convention sugar diet.

**Figure 6.**
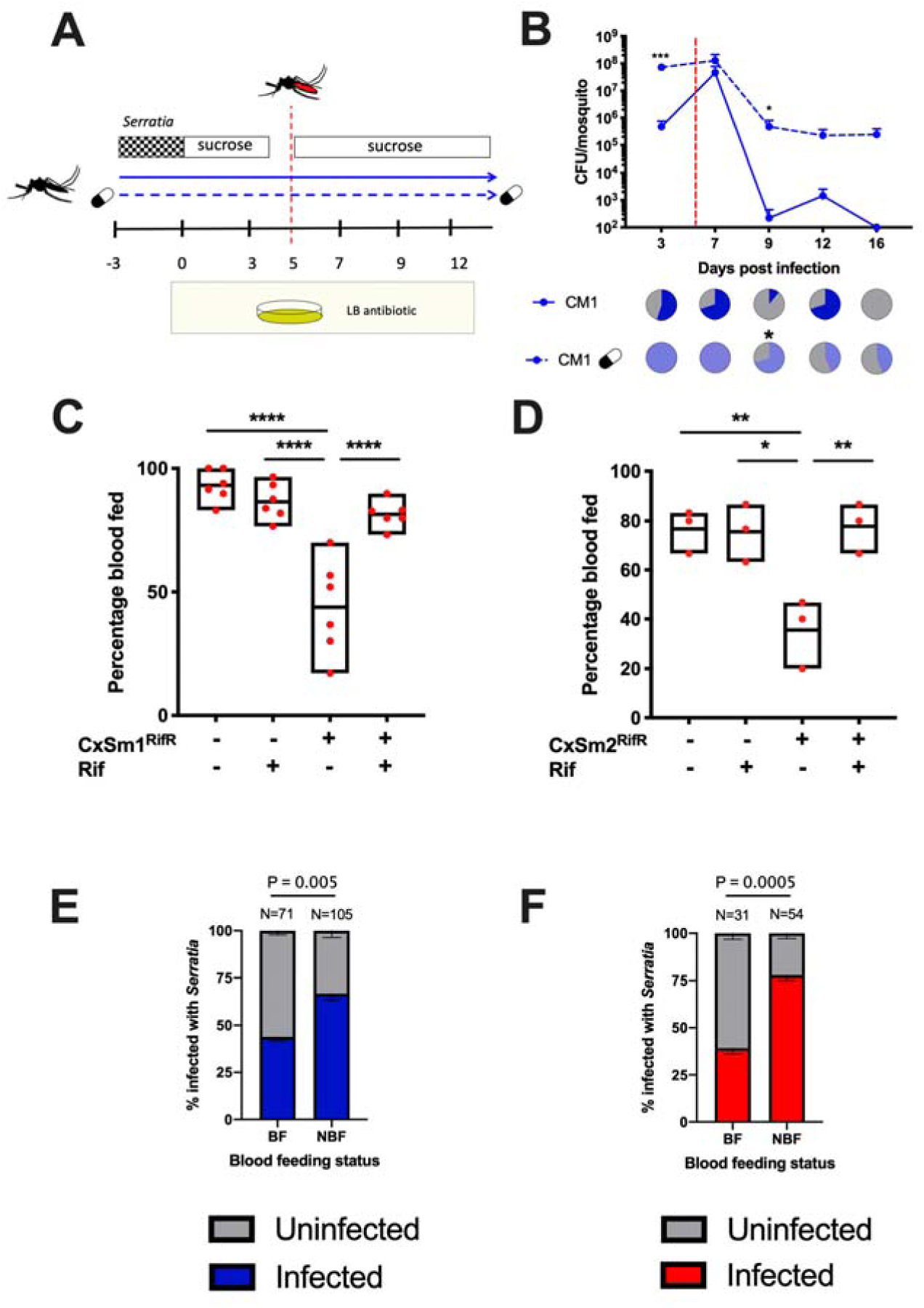
Interaction between *Serratia* infection and blood feeding in *Ae. aegypti*. Schematic depicting the infection and blood feeding experimental design (**A**). Infection density and prevalence of CxSm1^RifR^ in conventional and antibiotic fed *Ae. aegypti* (**B**). The red dotted line indicates the timing of blood meal. Significance was determined using a T-test comparing conventional and antibiotic groups for each time point. Ten mosquitoes were examined at each time point. Percentage of mosquitoes to take a blood meal for CxSm1^RifR^ (**C**) or CxSm2^SmR^ (**D**) infected or uninfected mosquitoes. Significance was determined using a one-way ANOVA with Tukey’s multiple comparisons test, * p <0.05, ** p < 0.01, **** p < 0.0001. Either six or three cups were used for feeding experiments with 50 mosquitoes per cup. Percentage *Serratia* administered mosquitoes infected with CxSm1^RifR^ (**E**) or CxSm2^SmR^ (**F**) blood fed, or non-blood fed groups. Fisher’s exact test was used to determine significance. Samples size is indicated for each group above the bars.

We, therefore, investigated whether *Serratia* infection altered mosquito blood feeding behaviour. After providing mosquitoes with the opportunity to feed, we saw significantly fewer females had imbibed a blood meal compared to uninfected or antibiotic treated CxSm1^RifR^ infected mosquitoes (**Fig 6C**, ANOVA p < 0.001). Blood-feeding rates in *Serratia* infected *Ae. aegypti* were restored when mosquitoes were fed antibiotics, indicating these behavioural changes were mediated by the interplay between CxSm1^RifR^ and other bacterial constituents of the microbiome susceptible to antibiotics. Given this intriguing finding, we repeated these experiments with the CxSm2^SmR^ isolate. Similar to findings with its close relative, the CxSm2^SmR^ *Serratia* strain altered the blood feeding rates in mosquitoes (Fig 6D, ANOVA p < 0.001). Given the heterogeneity in the prevalence of CxSm1^RifR^ and CxSm2^SmR^ in conventionally reared mosquitoes, we examined individuals that did or did not blood feed for *Serratia* infection. For both CxSm1^RifR^ (**Fig 6E**) and CxSm2^SmR^ (**Fig 6F**), the *Serratia* infection rate was significantly higher in non-blood fed mosquitoes compared to blood fed (CxSm1^RifR^ p < 0.005; CxSm2^SmR^ p < 0.005), indicating that mosquitoes that took a blood meal were less likely to be infected with *Serratia*. When considering this, it is likely the reductions we observed at the population level (**Fig 6C & D**) are conservative, and the effect of *Serratia* infection on blood feeding behaviour is more pronounced.

## Discussion

The interplay between the host and microbes can dictate insect microbiome homeostasis, but little is known regarding how microbe-microbe interactions within the gut influence microbial composition and abundance. Previously we identified a *Serratia* infection gradient in the arboviral vectors, *Ae. aegypti, Ae. albopcitus*, and *Cx. quinquefasciatus*, with high loads in the latter and an absence of infection in the former^4^. Here we show that *Serratia* poorly infects many *Ae. aegypti* strains and that the mechanism mediating this incompatibility is competitive exclusion from other members of the *Enterobacteriaceae*, which are close relatives of *Serratia.* Given that *Serratia* can influence vector competence in mosquitoes and has been proposed as a microbe for paratransgenic control^66,67^, it is imperative we enhance our understanding regarding the factors that influence *Serratia* acquisition in the mosquito gut.

After confirming that microbiota was inhibiting *Serratia* colonization of mosquitoes, we characterized the microbiome of *Ae. aegypti* lines susceptible and resistant to infection. Intriguingly, the susceptible Thailand line possessed a distinct and species-rich microbiome, and had significantly lower levels of *Enterobacteriaceae*. We speculate that this line had lost its capacity to maintain microbiome homeostasis, which subsequently enabled numerous other bacterial species to colonize. These other species likely reduce the abundance of *Enterobacteriaceae* in the host, as our qPCR data indicated that the Thailand line has similar bacterial load compare to the other lines. The reduced levels of antagonistic *Enterobacteriaceae* in the Thailand line enabled the colonization of the *Serratia* at levels similar to *Culex* mosquitoes. This theory is further supported by the fact that inhibition of *Serratia* was restored in the Thailand line when mosquitoes were pre-infected with *Cedecea*.

Microbiome dysbiosis can profoundly alter several host phenotypes in insects, including symbiont processes^18^. In the Oriental fruit fly, *Bactrocera dorsalis*, suppression of the dual oxidase gene (BdDuox) led to microbiome dysbiosis and an over-abundance of *Verrucomicrobiaceae* bacteria^69^. Increases in V*errucomicrobiaceae* have also been observed in mammalian systems when the microbiome transfers to a dysbiotic state^70-73^. In our analysis, *Verrucomicrobiaceae* was a dominant member of the microbiome of the Thailand line, yet it was at relatively low abundance in the Iquitos and Juchitan lines and barely detectable in the Galveston line. The presence of this family indicates the microbiome of the Thailand line was in a state of dysbiosis. In the Galveston line, the co-infection experiments recapitulated our previous results indicating bacterial co-exclusion was the main factor driving *Serratia* incompatibility. However, the effects in the Thailand line were more subtle, with the presence of *Cedecea* only reducing the *Serratia* infection prevalence at earlier time points and not influencing titer. This suggests that other host factors likely contribute to the incompatibility of *Serratia* in the Galveston line, but these factors were deficient in the Thailand line, which resulted in the more subtle phenotype. In *Galleria mellonella*, the greater wax moth, both host and bacterial factors synergize to control microbiome composition. When host immunity is suppressed, and mutant symbionts that lack the capacity to produce bacteriocins (proteins that inhibit closely related bacterial strains) are administered to the moth, *Serratia* proliferates within the microbiome^74^. In *Anopheles gambiae*, mosquitoes regulate *Serratia* infections by the complement pathway, and silencing of CLIP genes increases *Serratia* load, which subsequently induced mortality^75^. Similar to the Oriental fruit fly, in mosquitoes, Duox maintains redox homeostasis which in turn regulates microbiota^13,76^. Taken together, these studies suggest host and microbial factors together maintain microbiome homeostasis in mosquitoes, and when this is disrupted, other taxa that would normally be excluded can proliferate within the microbiome.

*Serratia* has pathogenic effects in several insect species^74,77-79^, but this bacterium is also a common taxa within the insect microbiome^23-28^. In hematophagous arthropods, there is variation in *Serratia’s* pathogenicity, ranging from inducing mortality or severe fitness costs on the host under certain conditions to having no observed effect^36,78,80^. We saw little evidence for *Serratia* affecting mortality or reproduction, similar that observed in *Culex* mosquitoes^81^. However, intriguingly, our data suggests that *Serratia* can altered the propensity of mosquitoes to take a blood meal.

While there are several examples that a broad range of microbes can influence feeding behavior in insects, relatively little is known regarding how gut-associated microbes contribute to these phenotypes. In flies, microbial communities affect affect feeding preference and egg laying behaviour^82,83^, and in mosquitoes, pathogens can alter feeding behaviour. For example, the fungus *Metarhizium* reduces blood feeding rates in *An. gambaie*^84^, while arbovirus infections in *Aedes* mosquitoes can alter feeding phenotypes^85,86^, potentially by altering expression of odorant binding proteins^87^. Alternatively, in several insect systems microbe-mediated alteration in immunity affects feeding behaviour^88-90^. *Plasmodium* infection in *Anopheles* alters host-seeking response, but similar phenotypes are also induced by microinjection of heat killed *Escherichia coli*, indicating immune challenge may mediate these behavioural phenotypes^91^. In *An. gambiae* there is an interplay between *S. marcescens* and gustatory receptors and odorant binding proteins^92^, and in flies, these gustatory receptors have been implicated in influencing behaviour^93,94^. Our data indicate that *Serratia* acts in concert with other microbes to reduce blood feeding. There is a complex immune interplay between gut microbes and the host^95-97^, and it is possible that disruption of microbiome homeostatis by *Serratia* infection may alter basal immunity which subsequently affects feeding behaviour. Alternatively, these mosquitoes may be suffering the effects of infection or microbiome dysbiosis resulting in a lack of interest in feeding.

From a vector control standpoint, reducing blood-feeding rates will greatly influence pathogen transmission. However, this phenotype is mediated by an interaction between *Serratia* and other native microbes of the *Ae. aegpyti*. Given the inherent variability in the microbiome of mosquitoes, further investigations are warranted to determine how universal this phenotype is, and in general how microbiome dysbiosis alters mosquito behaviour that can impact vectorial capacity. In the laboratory setting, reduced feeding rates would act as a distinct mechanism to eliminate *Serratia* infections from the microbiome of *Ae. aegypti*.

Another important aspect of our work is the finding that *Ae. aegpyti* lines reared under uniform insectary conditions have diverse microbiomes. While it is clear the Thailand line has a particularly divergent microbiome, the microbiomes of the Juchitan, Galveston, and Iquitos lines were also distinct from each other. This is contrary to a recent finding^39^, and suggests that similarity in microbiomes driven by environmental factors are not universal, and host or bacterial factors also play a role in microbiota community assembly and can lead to microbiome divergence. A recent analysis of diverse *Drosophila* species, reared under uniform laboratory conditions, found the composition of the microbiota varied^98^ while genotypically divergent *D. melanogaster* lines influenced commensal bacterial levels when reared under mono-axenic gnotobiotic conditions^99^. Here we demonstrate in mosquitoes that host genotype profoundly alters bacterial microbiome composition.

In conclusion, we show that microbe-microbe interactions influence microbiome composition and abundance in mosquito vectors. These processes are robust and can prevent the transfer of microbiota between mosquitoes that share a common environment by distinct mechanisms. Transfer of microbiota can occur in a host when microbiome homeostasis is disrupted, but this can also alter phenotypes important for host biology. Furthermore, we show that microbiota transfer can change mosquito traits that are important for pathogen transmission. From an applied standpoint, a greater understanding of the factors dictating microbial exclusion and acquisition could be exploited to develop strategies to create mosquitoes with designer microbiomes that induce desirable properties for vector control.

## Acknowledgements

We would like to thank the UTMB insectary core for providing mosquitoes and Alvaro Acosta-Serrano for commenting on a previous draft. GLH was supported by the BBSRC (BB/T001240/1), the Royal Society Wolfson Fellowship (RSWF\R1\180013), NIH grants (R21AI124452 and R21AI129507), the Western Gulf Center of Excellence for Vector-borne Diseases (CDC grant CK17-005) and the NIHR (NIHR2000907). GLH is affiliated to the National Institute for Health Research Health Protection Research Unit (NIHR HPRU) in Emerging and Zoonotic Infections at University of Liverpool in partnership with Public Health England (PHE), in collaboration with Liverpool School of Tropical Medicine and the University of Oxford. GLH is based at LSTM. The views expressed are those of the author(s) and not necessarily those of the NHS, the NIHR, the Department of Health or Public Health England. This work was also supported by a James W. McLaughlin postdoctoral fellowship at the University of Texas Medical Branch to SH, and a NIH T32 fellowship (2T32AI007526) to MAS.

## Supplementary figures and files

**Figure S1. Phenotypic characterisation of Serratia isolates.** Swimming motility of CxSm1 and CxSm2 at different temperatures (**A**). Oxidase activity at 30°C (**B**). Scanning electron microscopy of CxSm1, CxSm2, and *Cedecea*.

**Figures S2. Genomic analysis of CxSm1 and CxSm2 strains.** Our isolates were compared to a set of publicly available *Serratia* reference genomes (Table S3) using average nucleotide identity (ANI). The heatmap indicates pairwise comparisons of ANI in an all-against-all comparison, showing our isolates as a distinct subgroup within *S. marcescens*. Colour legends for the similarity (ANI) and the species designation as given in GenBank are detailed below (Table S3).

**Figure S3. Total culturable bacterial loads in mosquitoes.** Homogenized mosquitoes were plated on LB agar plates without selection. Line graph indicates bacterial density (CFU/mosquito) and pie graph indicated prevalence. Data related to Figure 1A.

**Figure S4. Native *Serratia* densities in *Cx. quinquefasciatus* strains.** qPCR relative abundance (*Serratia*/S7) values of the Houston, Florida and Salvador *Cx. quinquefasciatus* lines reared under identical conditions in the same insectary. Data were analysed with an ANOVA with Tukeys multiple comparison. (** p < 0.01, *** p < 0.001)

**Figure S5. Validation metrics of microbiome sequencing data.** Shannon entropy rarefied at intervals between 0 and 80000 reads for each individual (A). Relative adundance of positive spike-in controls (B).

**Figure S6. Microbiome relative abundance measures for each individual at the genus levels.**

**Figure S7. Survival curves of *Serratia* infected mosquitoes. Curves** relate to Figure 1A (**A**), Figure 4 (**B**), Figure 5 (**C**), and Figure 6 (**D**).

**Figure S8. Reproductive output of blood fed mosquitoes measured in terms of egg number.**

**Table S1. List of mosquito lines used in experiments.**

**Table S2. Primers used for PCR and qPCR.**

**Table S3. Selection of reference genomes used in comparative genomics.**

**Table S4: Average ANI distances of CxSm1 and CxSm2 against a selection of reference genomes (See Table S3); average values per species are shown.**

**Table S5. OTU table for 16S rRNA amplicon sequencing.**

