## Supplementary figures and images for "Microbial interactions in the mosquito gut determine *Serratia* colonization and blood feeding propensity"

### Fig S1

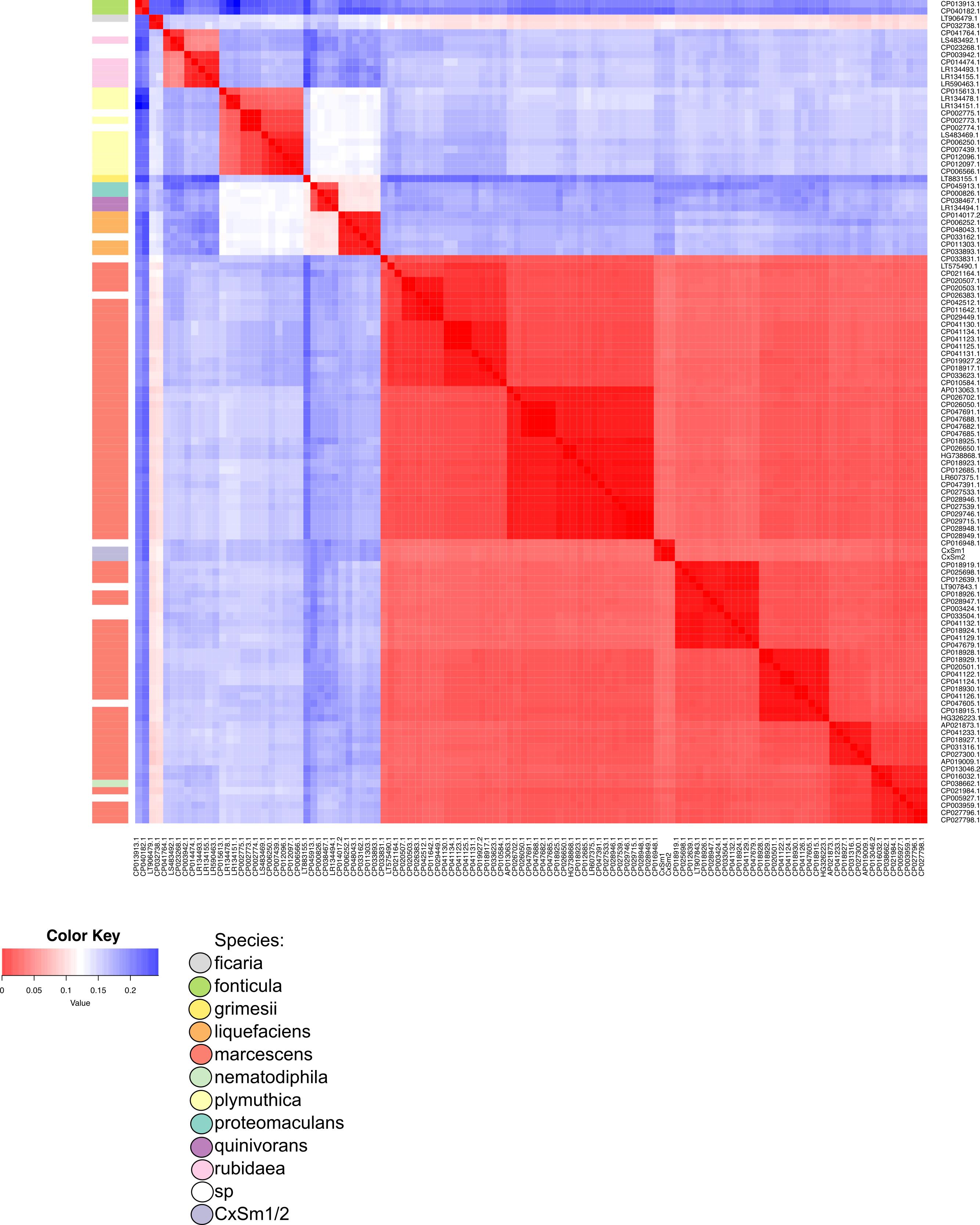

### Fig S2

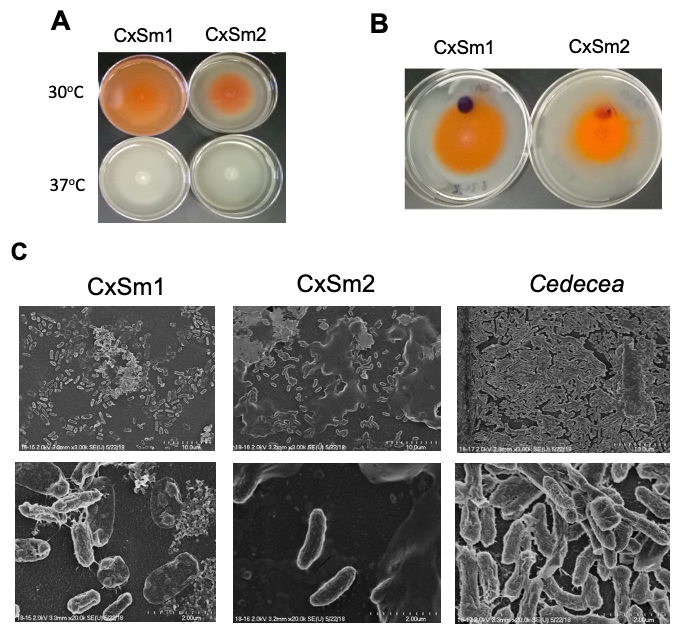

### Fig S3

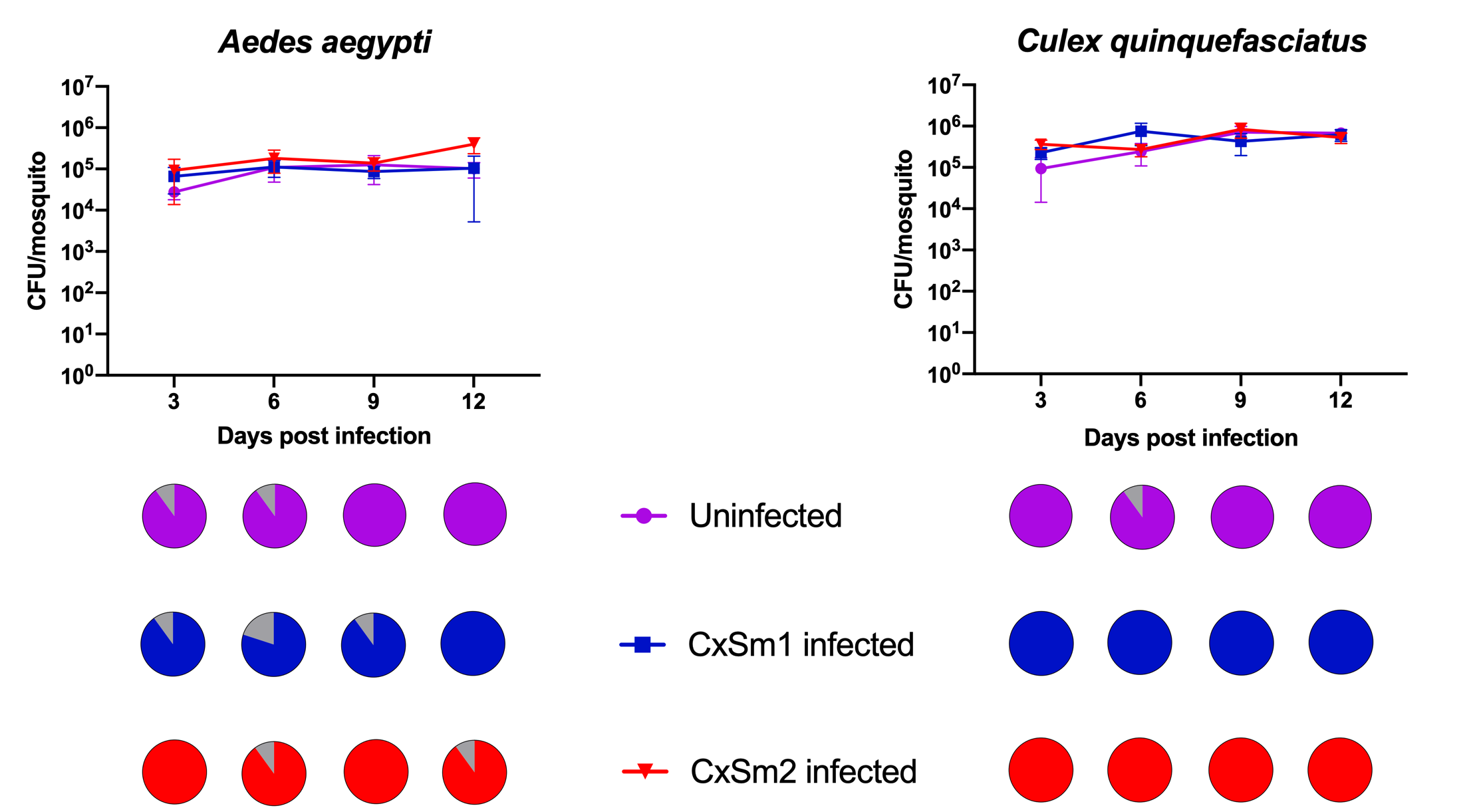

### Fig S4

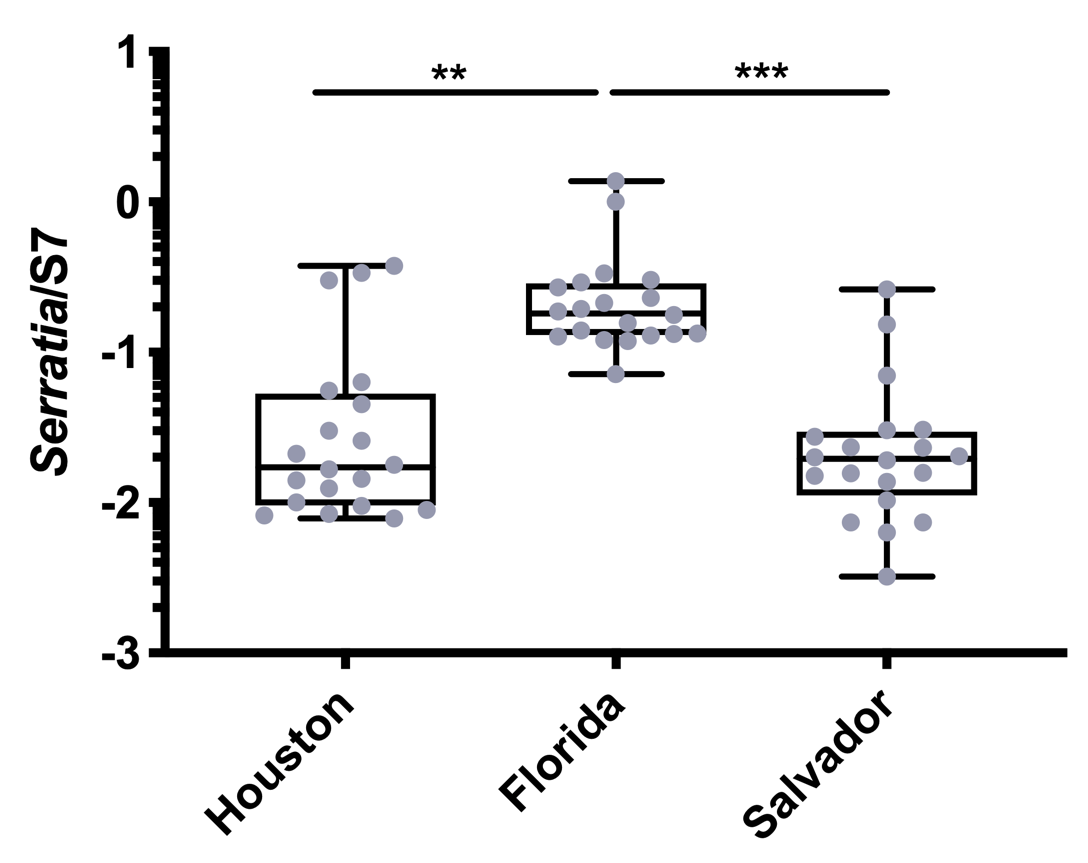

### Fig S5

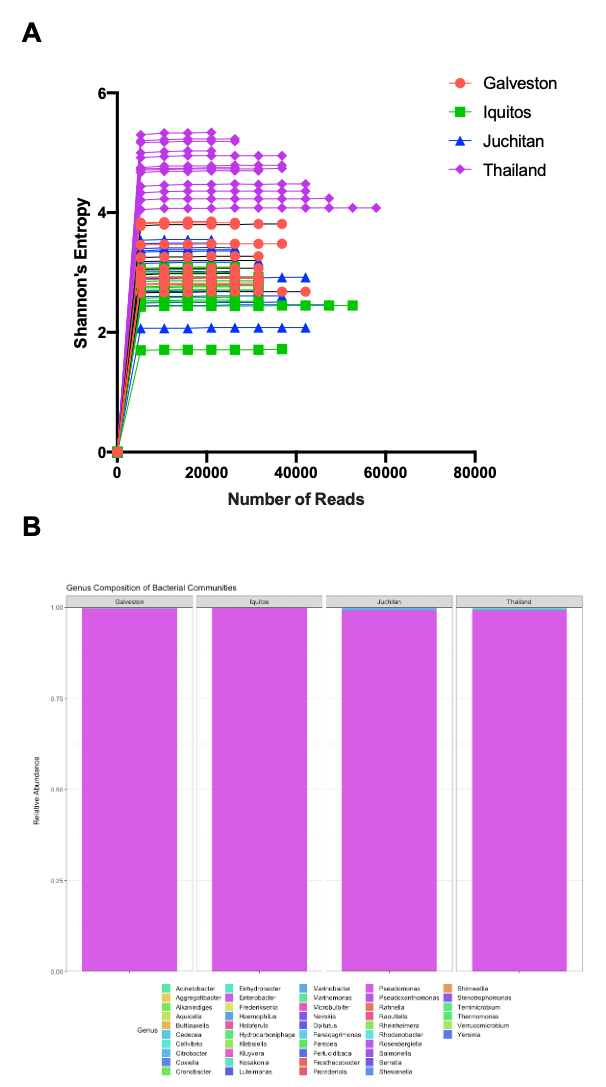

### Fig S6

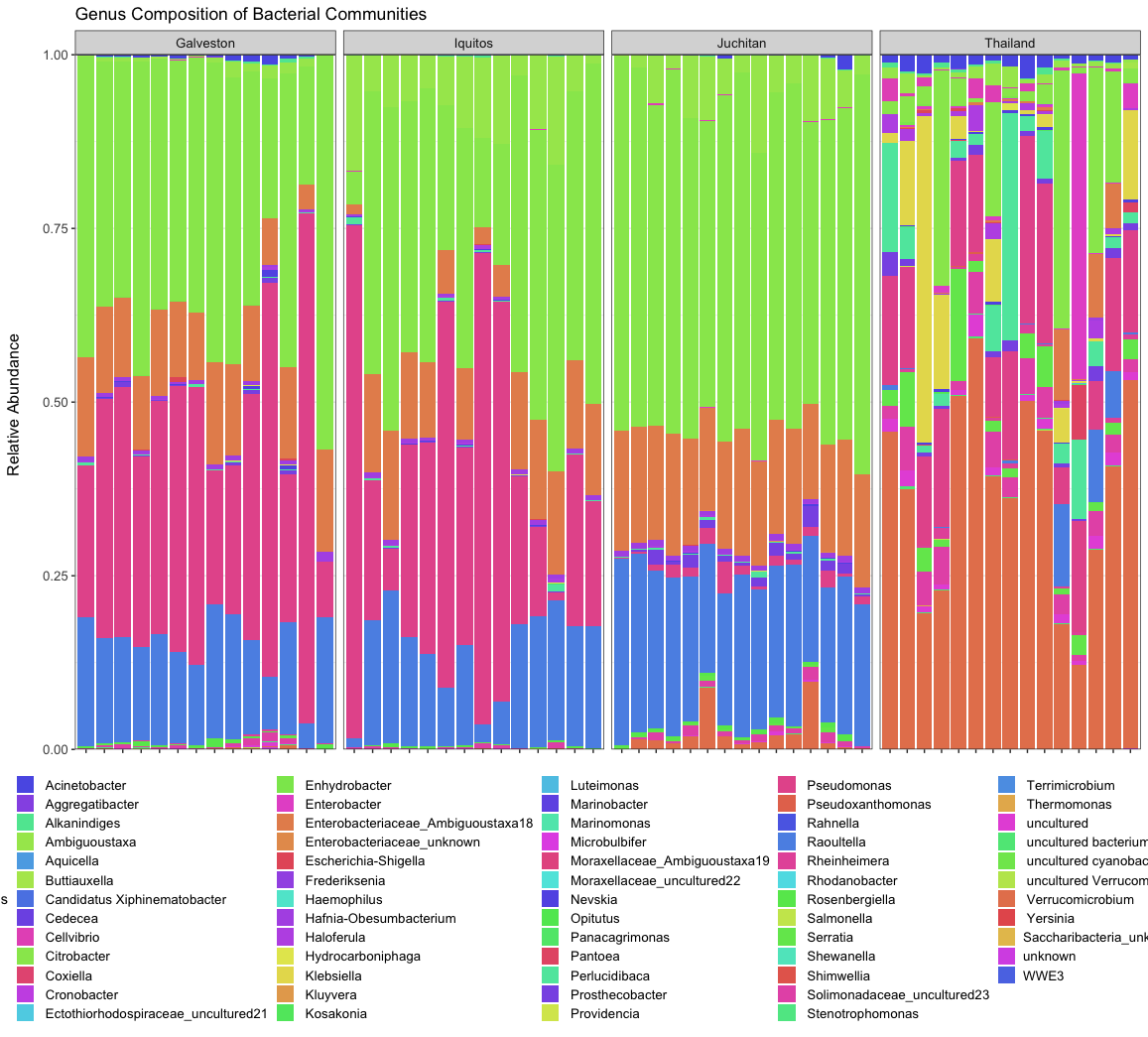

### Fig S7

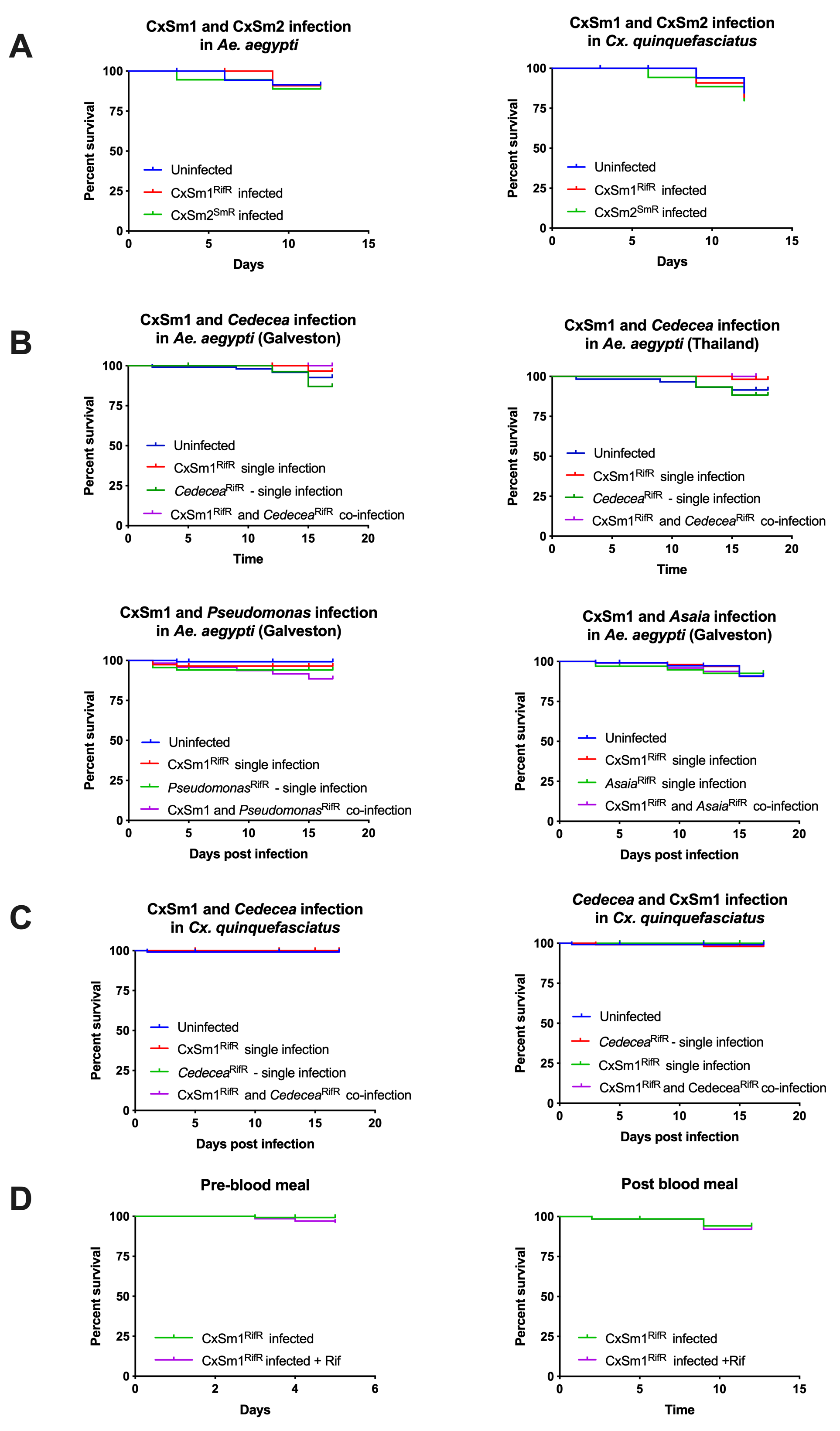

### Fig S8

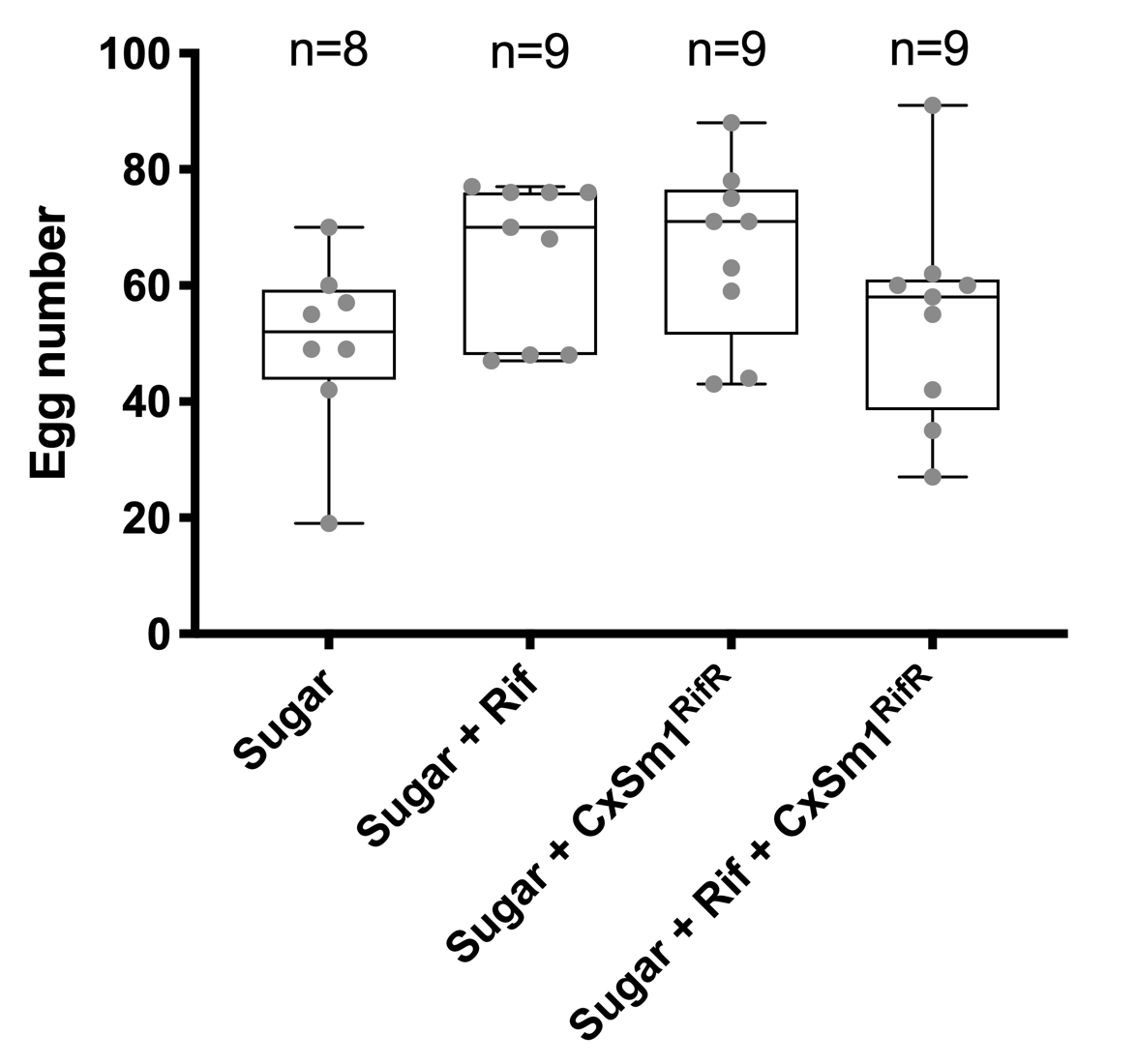
